## Supplemental Figures for "Time of day analysis over a field grown developmental time course in rice"

### Supplemental Figures and Tables

#### Tables

Table S1. Rice genes that are not expressed (le) or not rhythmic (nr).
Table S2. Genes that are expressed by not rhythmic across any of the time courses.
Table S3. Genes that cycle under all conditions including root.
Table S4. Genes that cycle in all leaf conditions.
Table S5. Core circadian clock, light and flowering genes.
Table S6. Significant cis-elements identified across the time courses.
Table S7. Sequence of significant cis-elements.
Table S8. Significant gene ontology (GO) terms.
Table S9. Significant GO terms by developmental time point with more than two occurrences over the day.
Table S10. TOD significance for cis-elements.

### Figures

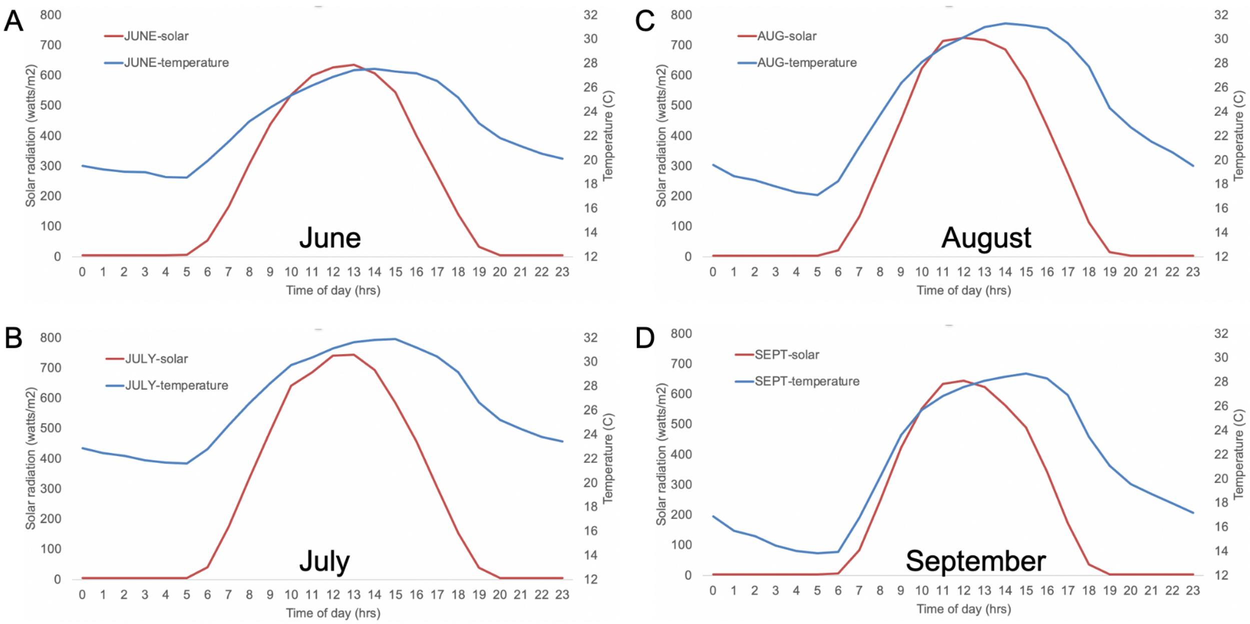


**Figure S1. Temperature lags daily changes in solar radiation.** The average solar radiation (red) was plotted against the average temperature for (A) June; (B) July; (C) August; (D) September. The values were generated by averaging the solar radiation or temperature by time of day (TOD).

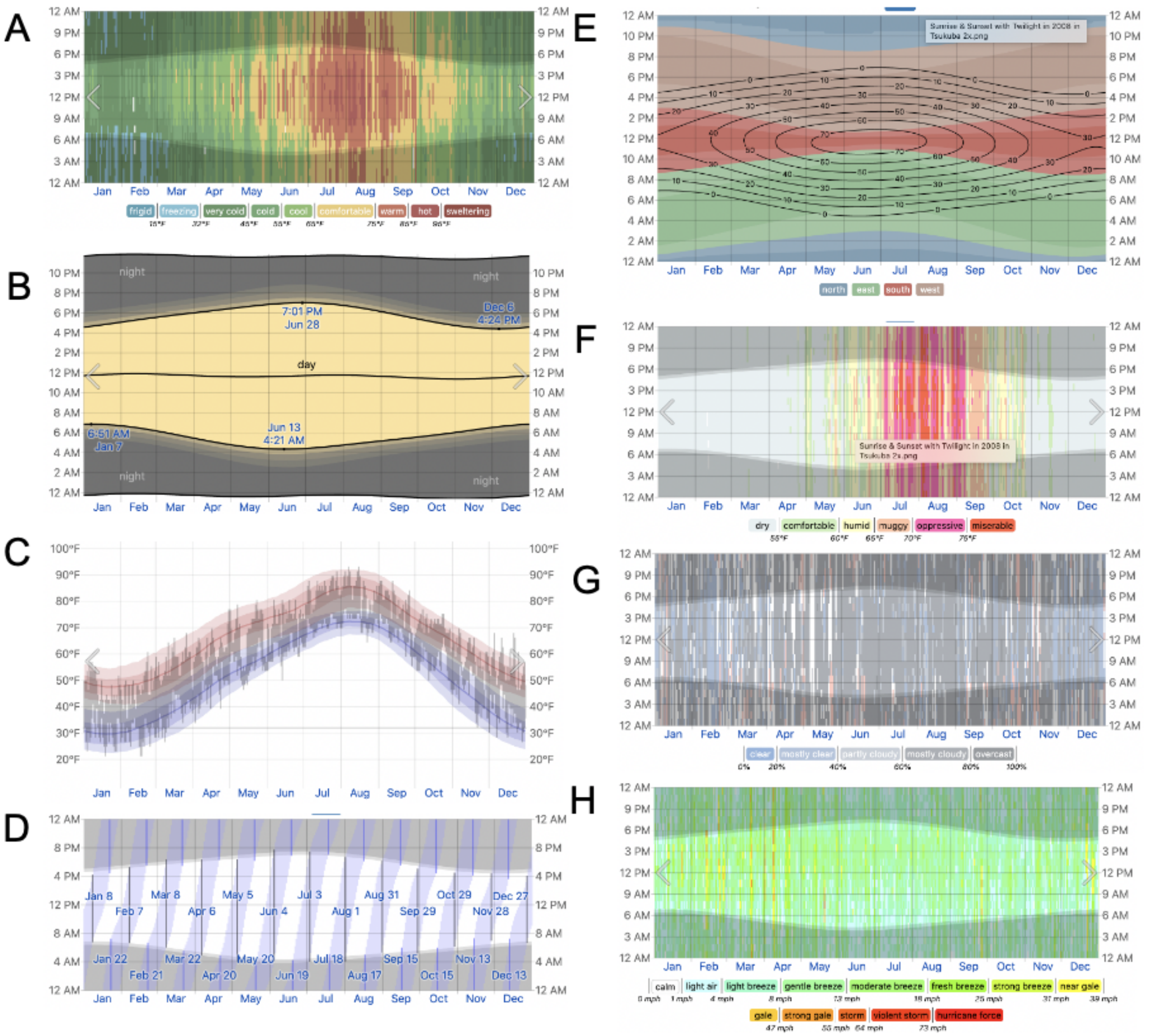


**Figure S2. Weather conditions in Tsukuba, Japan in 2008.** (A) Temperature cycles; (B) light/dark cycles/photoperiod; (C) average temperature; (D) moon cycle; (E) Solar Elevation and Azimuth; (F) humidity; (G) cloud cover; (H) wind speed. Data was downloaded from: weatherspark.com/h/y/144004/2008/Historical-Weather-during-2008-in-Tsukuba-Japan#Figures-ObservedWeather.

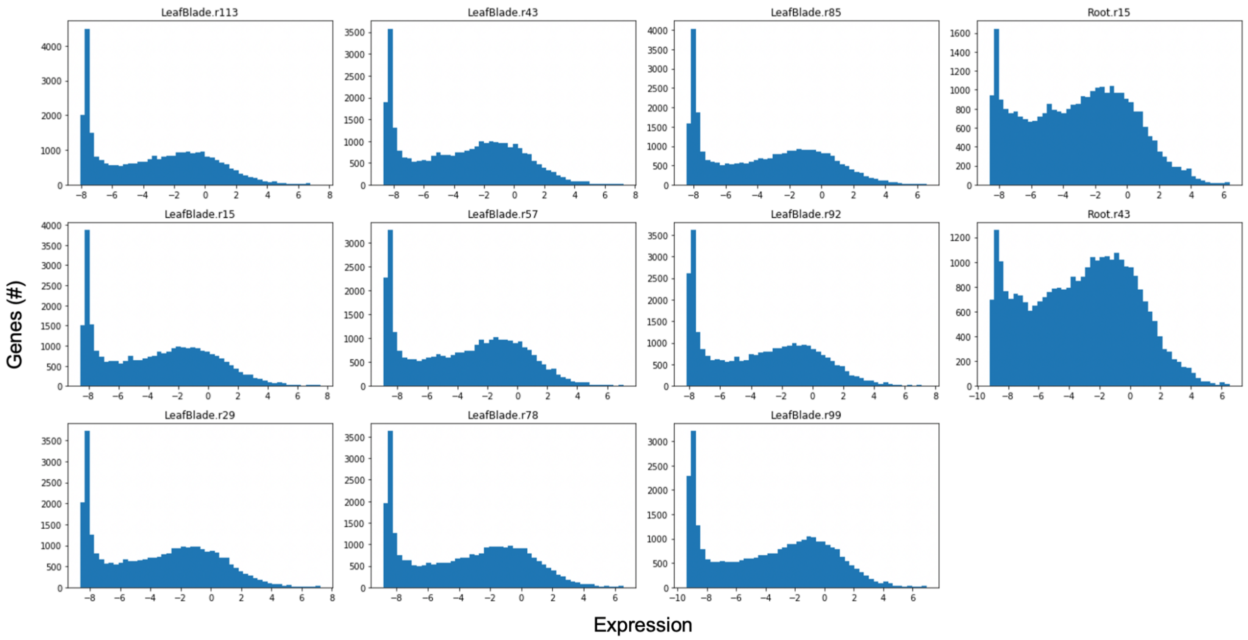


**Figure S3. Expression analysis to determine the cutoff for "non-expressed" genes.** Label above each plot for all eleven time courses. Expression level by number of genes is plotted to identify potential non-expressed genes.

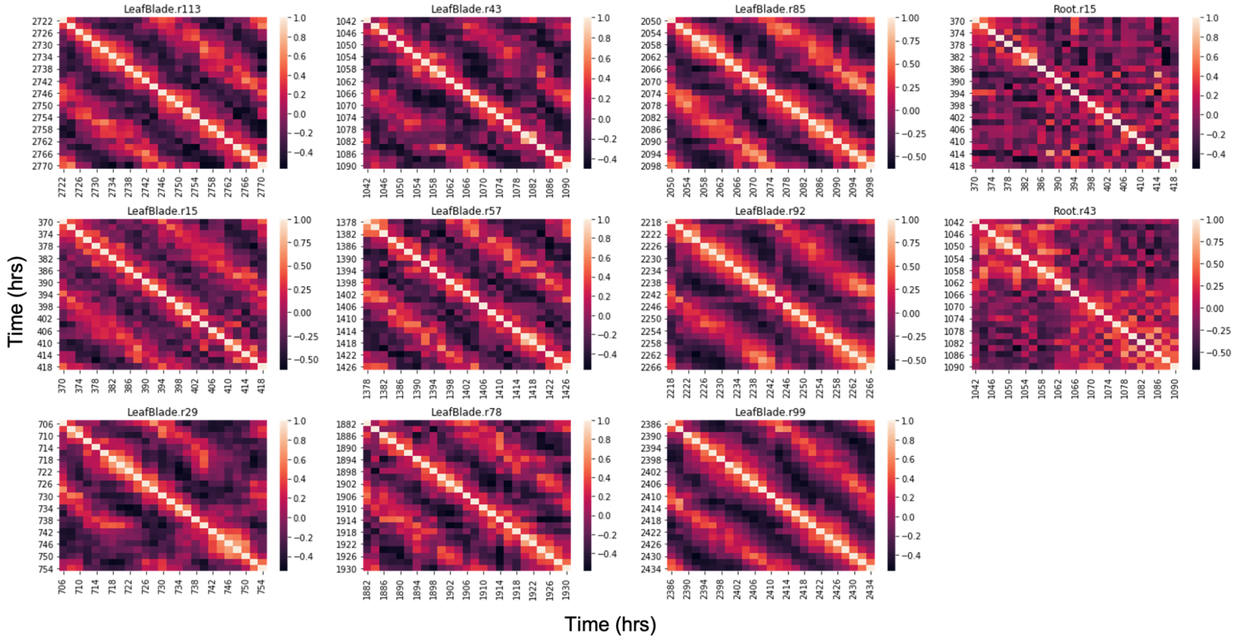


**Figure S4. Correlation of expression in each time course reveals patterns of cycling behavior.** Label above each plot for all eleven time courses. Time courses are organized by time and internal correlation shows the clear cycling behavior in the leaf time courses and the loss of coherent cycling in the two root time courses.

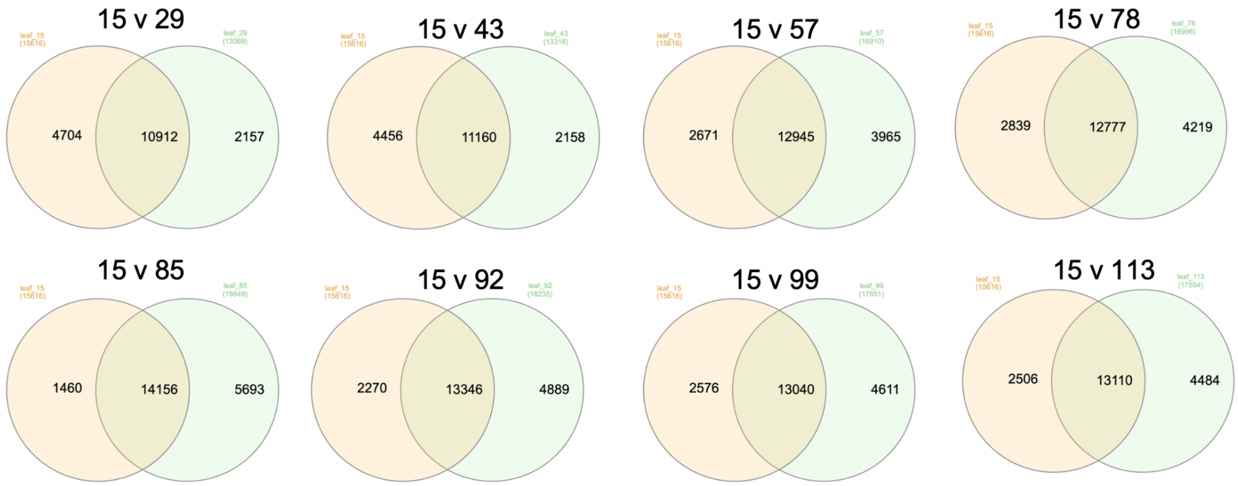


**Figure S5. Overlap of genes predicted to cycle compared to the first time course.** Genes that are predicted to cycle (R>0.8) were compared between time courses denoted above the venn diagram.

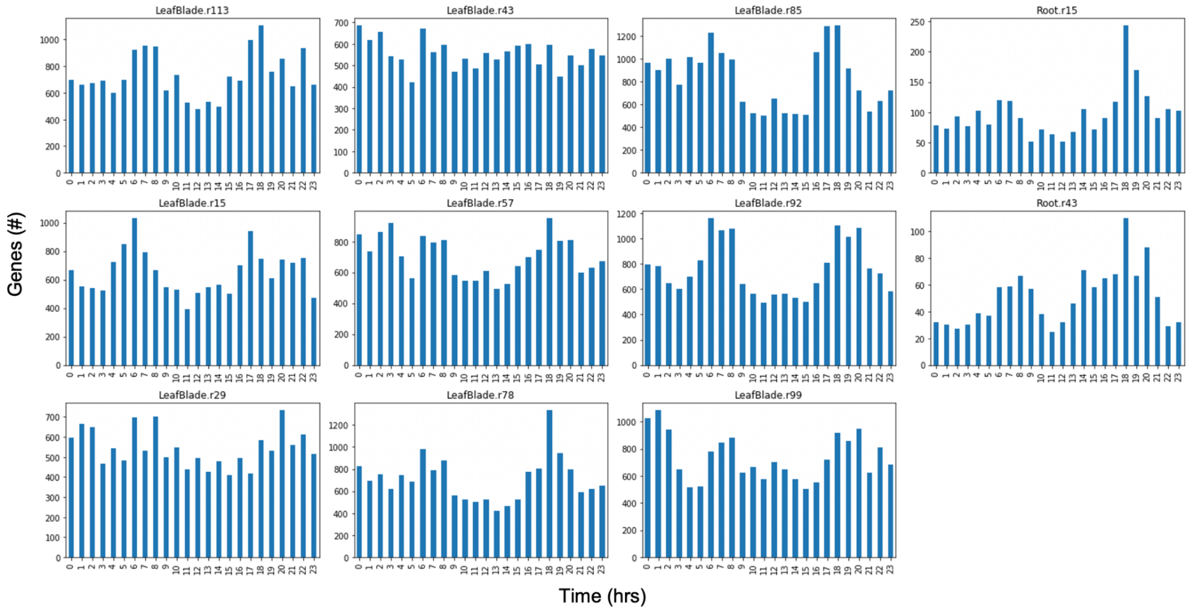


**Figure S6. Number of cyclers by time of day (TOD).** Label above each plot for all eleven time courses. Histogram of number of genes predicted to cycle at a specific TOD.

A

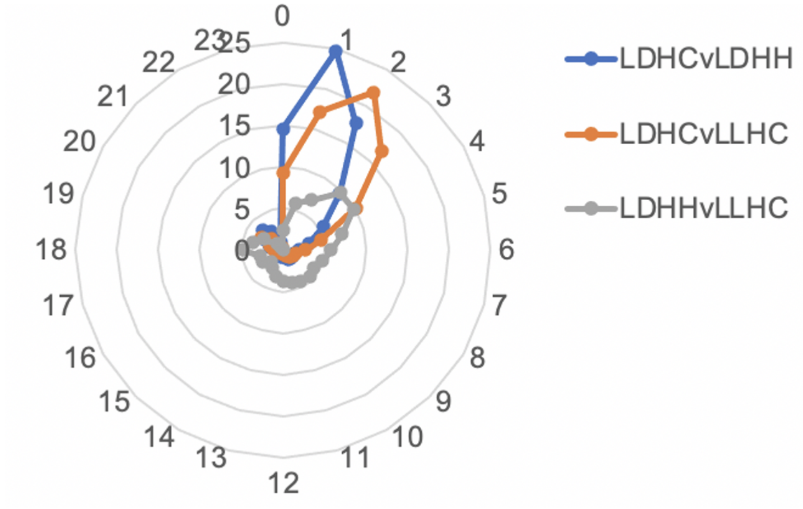


B

| | LDHCvLDHH | LDHCvLLHC | LDHHcLLHC |
|---|---|---|---|
| 0 | 15 | 9 | 2 |
| 1 | 25 | 17 | 6 |
| 2 | 18 | 22 | 7 |
| 3 | 10 | 17 | 10 |
| 4 | 6 | 10 | 10 |
| 5 | 3 | 5 | 7 |
| 6 | 2 | 3 | 6 |
| 7 | 1 | 2 | 5 |
| 8 | 1 | 1 | 4 |
| 9 | 1 | 1 | 5 |
| 10 | 1 | 1 | 4 |
| 11 | 1 | 1 | 4 |
| 12 | 1 | 0 | 4 |
| 13 | 1 | 0 | 3 |
| 14 | 0 | 0 | 2 |
| 15 | 1 | 0 | 2 |
| 16 | 1 | 0 | 3 |
| 17 | 1 | 1 | 3 |
| 18 | 1 | 1 | 5 |
| 19 | 2 | 2 | 4 |
| 20 | 2 | 3 | 3 |
| 21 | 3 | 2 | 1 |
| 22 | 3 | 1 | 0 |
| 23 | 1 | 0 | 0 |

**Figure S7. Light and temperature have differential impact on TOD expression in rice.** Data from Filichkin et al., was analyzed for the impact of light cycles only (light/dark, continuous hot; LDHH), temperature cycles alone (continuous light, hot/cold; LLHC), and both light and temperature cycles (LDHC). (a) Number of genes showing a phase shift between the conditions LDHC and LDHH (blue), LDHC and LLHC (orange), and LDHH and LLHC (gray). Time is on the circumference and number of genes on the radius; (b) percent of genes by the phase shift time (hrs) between the conditions. Green, high percentage; red, low percentage.

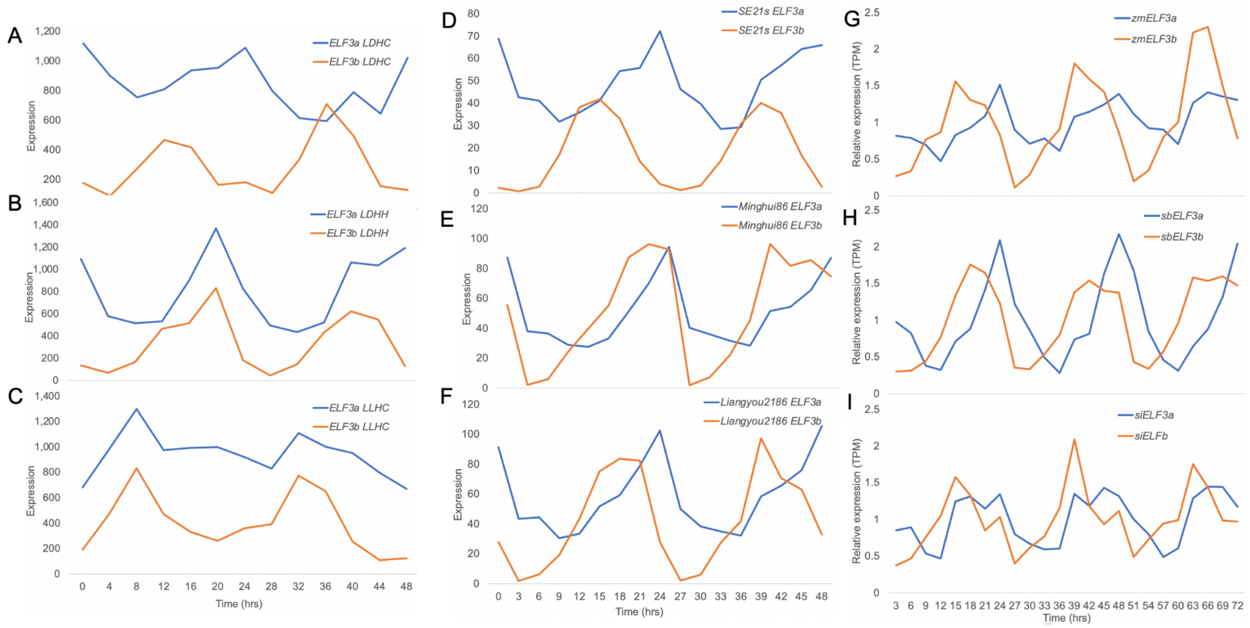


**Figure S8. ELF3 paralogs are responsive to light and temperature across grasses.** *ELF3a/ELF3-1/HD17* (blue) and *ELF3b/ELF3-2* (orange) were evaluated in three different datasets for the impact of light and temperature entertainment in rice, cycling in inbreds and hybrids in rice, and cycling in *Zea mays* (zm), *Sorghum bicolor* (sb), and *Setaria italica* (si). (A) In wild type (wt) rice under LDHC the two paralogs are expressed antiphasic to one another; (B) wt rice under LDHH the two paralogs are in phase with dawn expression; (C) wt rice under LLHC the two paralogs are in phase with dusk expression; (D) the male-sterile line SE21s under LDHC the two paralogs are expressed anti-phase; (E) the restorer line Minghui86 under LDHC the two paralogs are in phase with expression at dawn; (F) super-hybrid rice Liangyou2186, which is a cross between SE21s and Minghui86, the two paralogs have anti-phasic expression; (G) *Zea mays* (zm) under LDHC the two paralogs have anti-phasic expression; (H) *Sorghum bicolor* (sb) under LDHC the two paralogs have anti-phasic expression; (I) *Setaria italica* (si) under LDHC the two paralogs have anti-phasic expression.

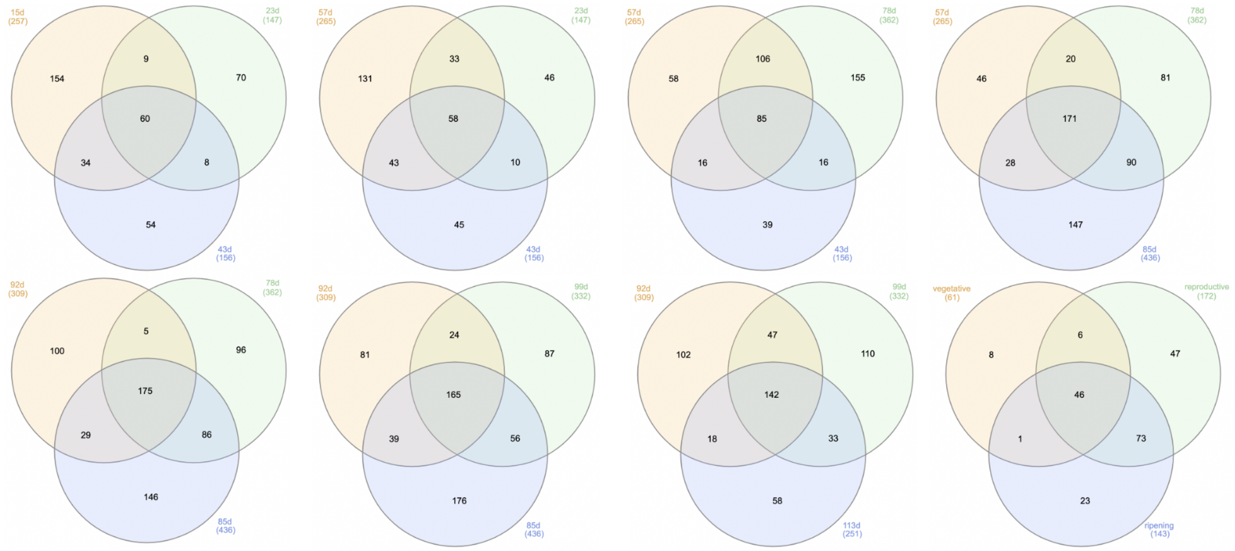


**Figure S9. Overlap of significant (P<0.05) gene ontology (GO) terms between conditions.** Conditions and number of GO terms used in the overlap denoted.

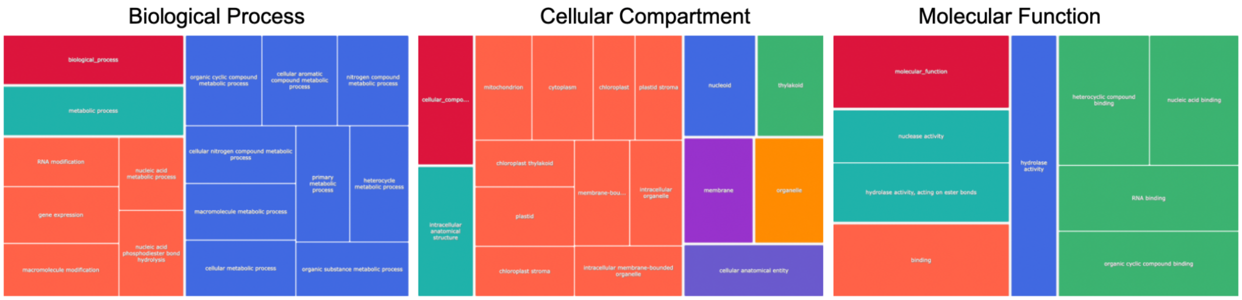


**Figure S10. Gene ontology (GO) term summary for the 46 shared terms across the nine leaf time courses.** The 46 GO terms were summarized using the REVIGO (REduce + VIsualize Gene Ontology) tools and plotted as Tree Maps for Biological Process (bp), Cellular Component (cc) and Molecular Function (mf).

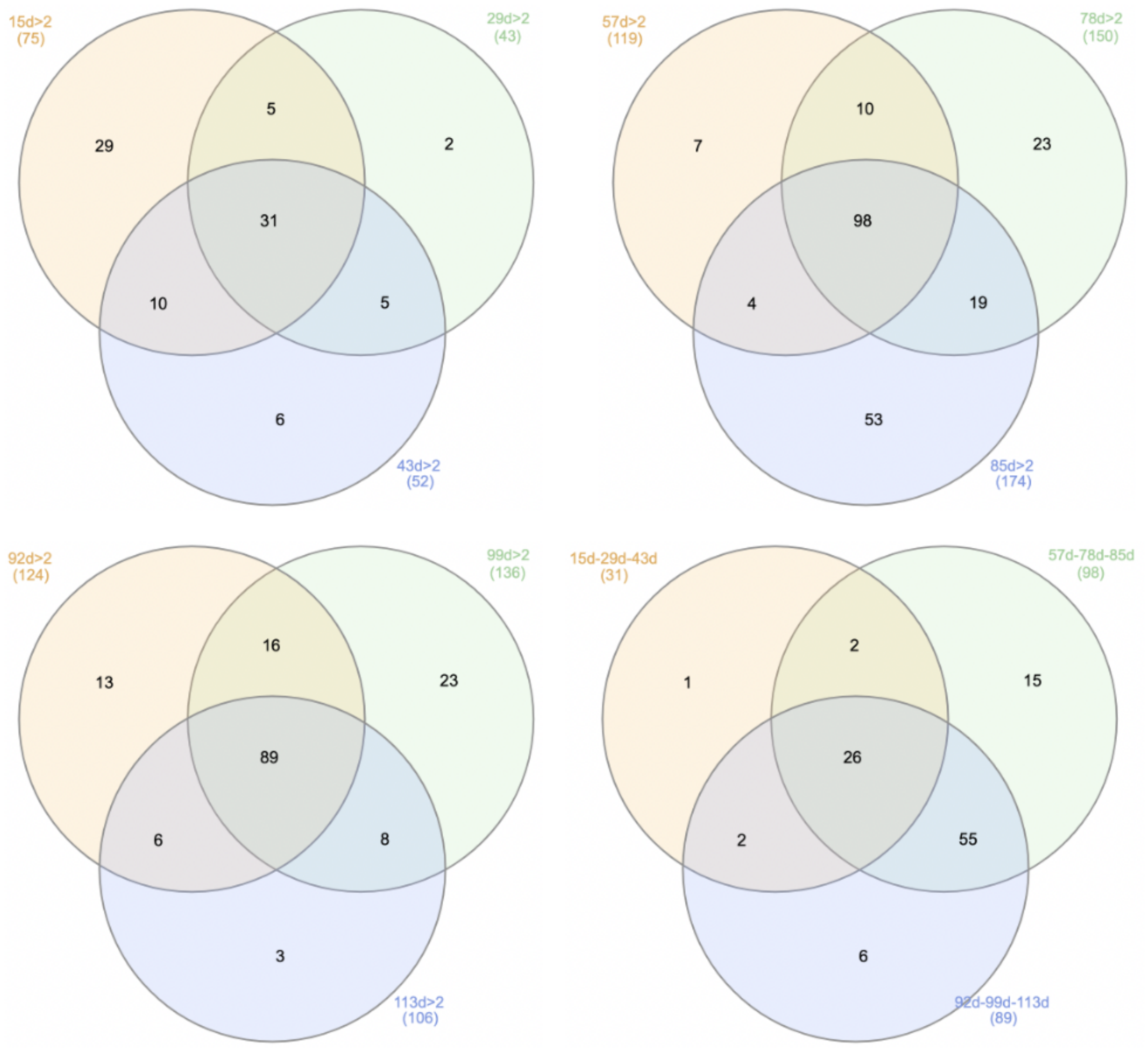


**Figure S11. Overlap of gene ontology (GO) terms that are significant under at least two times of day.** The condition plus the number of genes is denoted.
